# From stress to survival: identifying heat-resistant corals in Belize for conservation and restoration

**DOI:** 10.1101/2025.08.17.670758

**Authors:** DeVant’e D. Dawson, Nicole Craig, Courtney N. Klepac, Marilla Lippert, Brendan Cornwell, Wilbert Castillo, Calvin Quigley, Stephen R. Palumbi

**Affiliations:** Oceans Department, Hopkins Marine Station of Stanford University, Pacific Grove, United States of America; The Nature Conservancy, Belmopan, Belize; Department of Science and Technology, The University of Belize, Belmopan, Belize; Department of Geology and Geophysics, Woods Hole Oceanographic Institution, Woods Hole, United States of America

## Abstract

Rising ocean temperatures intensify coral bleaching, yet the response to high temperatures may also hold the key to reef survival. The potential lies in identifying individuals or populations that can withstand elevated temperatures, which could help inform more targeted conservation and restoration efforts. This study assessed the thermal resistance of four dominant coral species in Belize to identify populations best suited for long-term survival under increasing ocean temperatures. Coral populations from six characterized thermal reef sites were subjected to short-term heat exposure to determine bleaching thresholds (ED50), using Visual Bleaching Scores and experimental Degree Heating Days. As in other studies, we found a wide range of heat tolerance within and among species. The massive starlet coral (*Siderastrea siderea*) exhibited the highest resistance to thermal stress, while the brain coral *(Pseudodiploria strigosa)* was the most susceptible. The mountainous star coral (*Orbicella faveolata*) and mustard hill coral (*Porites astreoides*) showed intermediate heat resistance that varied by site. Notably, the shallower reefs (∼1-2 meters) of both inshore, central Belize and offshore Turneffe Atoll on the Belize Barrier Reef housed the highest proportion of the most thermally resistant corals. These findings highlight two key adaptive reef management opportunities: conserving naturally heat-resistant reefs and prioritizing thermally tolerant corals in transplantation efforts for restoration. By identifying corals and reef sites with the highest survival potential, this study provides essential insights to help safeguard Belize’s reefs against climate change, ensuring their continued ecological and economic benefits.

## Introduction

The Caribbean Sea has experienced rapid warming in recent decades, with sea surface temperatures increasing by up to 0.5℃ per decade in some areas [1]. This rate is more than twice the global average for the Northern Hemisphere, which was approximately 0.19 ± 0.13 °C per decade from 1979-2005 [1]. This warming, combined with other local and regional stressors, has contributed to widespread loss of hard coral cover across the region. On average, hard coral cover has declined by 80%, dropping from approximately 50% to 10% cover between the 1970s and early 2000s [2,3].

Belize, located within the Mesoamerican Barrier Reef System (MBRS), the second-largest reef system in the world, follows this trajectory. The Belize MBRS spans 190 miles and supports more than 60 coral species [4,5]. Between 2004 and 2014, maximum summer temperatures regularly exceeded the regional bleaching threshold of 31.5°C [6]. Some locations recorded over 14 consecutive days above this critical threshold, leading to severe bleaching and coral mortality [6]. Lagoon environments, in particular, experience even higher temperatures, with annual mean temperatures rising by 0.0805°C per year (∼ 0.8°C per decade) — a rate that significantly surpasses broader regional warming trends [6]. Consequently, hard coral cover across the Belize MBRS has declined by approximately 20% over the past two decades [7].

These warming trends and associated losses jeopardize not only biodiversity but also the livelihoods that depend on reef health. Belize’s coral reefs are vital to the nation’s ecological and economic well-being, supporting fisheries, tourism, and cultural traditions and contributing up to US$559 million annually to the economy [8]. Traditional reef management strategies, such as reducing pollution [9–12] and regulating overfishing [13–15], are essential for mitigating local stressors. However, these measures alone cannot counteract the accelerating pace of ocean warming, structurally altering coral reef ecosystems [16–19]. If current warming trends continue, corals in the Caribbean may experience annual bleaching within the next two to three decades, with yearly bleaching events projected as early as 2040 [20,21]. Without a corresponding increase in thermal tolerance of 0.2–1.0°C per decade, large-scale bleaching events in the Caribbean and beyond will persist [3].

To ensure the long-term survival of Belize’s reefs, conservation strategies must extend beyond traditional management and incorporate innovative approaches that enhance reef bleaching resistance [17–19,22]. One promising strategy is identifying and protecting heat-tolerant corals — species and populations that have naturally adapted to withstand extreme thermal conditions [23–31]. Experimental work has shown that some corals sourced from naturally warmer reef environments maintain higher bleaching resistance and survivorship under thermal stress, supporting their use in restoration efforts [32]. Research suggests that corals inhabiting thermally variable environments, possibly including Belize’s lagoons, may have greater resistance to rising temperatures [3,33]. These previous studies also show variability in heat resistance among corals within even the warmest reef environments, suggesting the need for testing heat resistance even at these sites.

Reef environments in Belize span a natural gradient of thermal conditions, with inshore and lagoonal sites experiencing significantly higher and more variable temperatures than offshore locations [3]. To test for variation in heat tolerance across this range of environmental variability, we selected sites across the Belize Barrier Reef Reserve System that represent a range of reef conditions. At Turneffe Atoll, sampling included a shallow reef crest site on the northeastern side, subject to open-ocean conditions and wave action [34], as well as sheltered sites along the atoll’s western side. We also sampled shallow, inshore reefs near the central Belize coastline, which experience limited water exchange and elevated thermal variability [3]. By assessing heat resistance within and between different coral species and reef environments, this study aims to develop methods for testing coral heat tolerance in Belize and identify reef populations that may serve as ecological strongholds for conservation and restoration initiatives.

Here, we present a multi-site assessment of heat tolerance variation across 206 coral colonies from four species across six sites within the Belize Barrier Reef Reserve System. These corals were subjected to standardized short-term heat stress assays to determine their bleaching thresholds (ED50), a metric of cumulative heat exposure at which corals appear half bleached (i.e., ∼50% loss of color or a Visual Bleaching Score (VBS) of 3). ED50 values were calculated using the linear regression of VBS, a rapid assessment of bleaching severity recorded during the experiment, against experimental Degree Heating Days (eDHD), which quantify cumulative thermal stress.

Our results reveal significant variation in thermal tolerance within and across species, as well as among locations, with some colonies demonstrating higher resistance to elevated temperatures. The stress-tolerant *S. siderea* exhibited high heat tolerance across sites. For other species, we found greater tolerance in a shallow, warm inshore reef and the shallow zone of an offshore atoll reef. These findings suggest that heat-tolerant corals are widely distributed and can be found by standardized testing across locations and species.

By identifying these resistant coral populations, this study provides a valuable tool and actionable data that is already informing national conservation planning. Specifically, our results support Belize’s goal of tripling fully protected coral reef areas from 7% to 20% by 2026 under the Resilient Bold Belize Initiative, which includes the Belize Sustainable Ocean Plan (BSOP) and Project for the Permanence (PFP). This work can help local communities more effectively protect their reefs and sustain the ecological and economic benefits they provide.

## Results

### Species differences

We first assessed how quickly each species began bleaching by examining changes in mean Visual Bleaching Score (VBS) across experimental degree heating days (eDHD). Our results reveal significant differences in bleaching rates among species (*p* = .00153; Fig 1), as well as variation in overall thermal tolerance within and across species and locations, with some colonies showing higher resistance to elevated temperatures. *Siderastrea siderea* was the slowest to bleach, reaching the VBS ≥ 3 threshold after 6.4 eDHD and bleaching significantly less than *O. faveolata* (*p* = 0.008), *Ps. strigosa* (*p* = 0.014), and marginally less than *Po. astreoides* (*p* = 0.053). In contrast, *Po. astreoides* and *O. faveolata* reached moderate bleaching after 4.7-5 eDHD, while *Ps. strigosa* was the most heat-susceptible, reaching the same bleaching threshold after only 3.4 eDHD. No significant differences were observed between *O. faveolata, Po. astreoides, and Ps. strigosa*.

**Fig 1.**
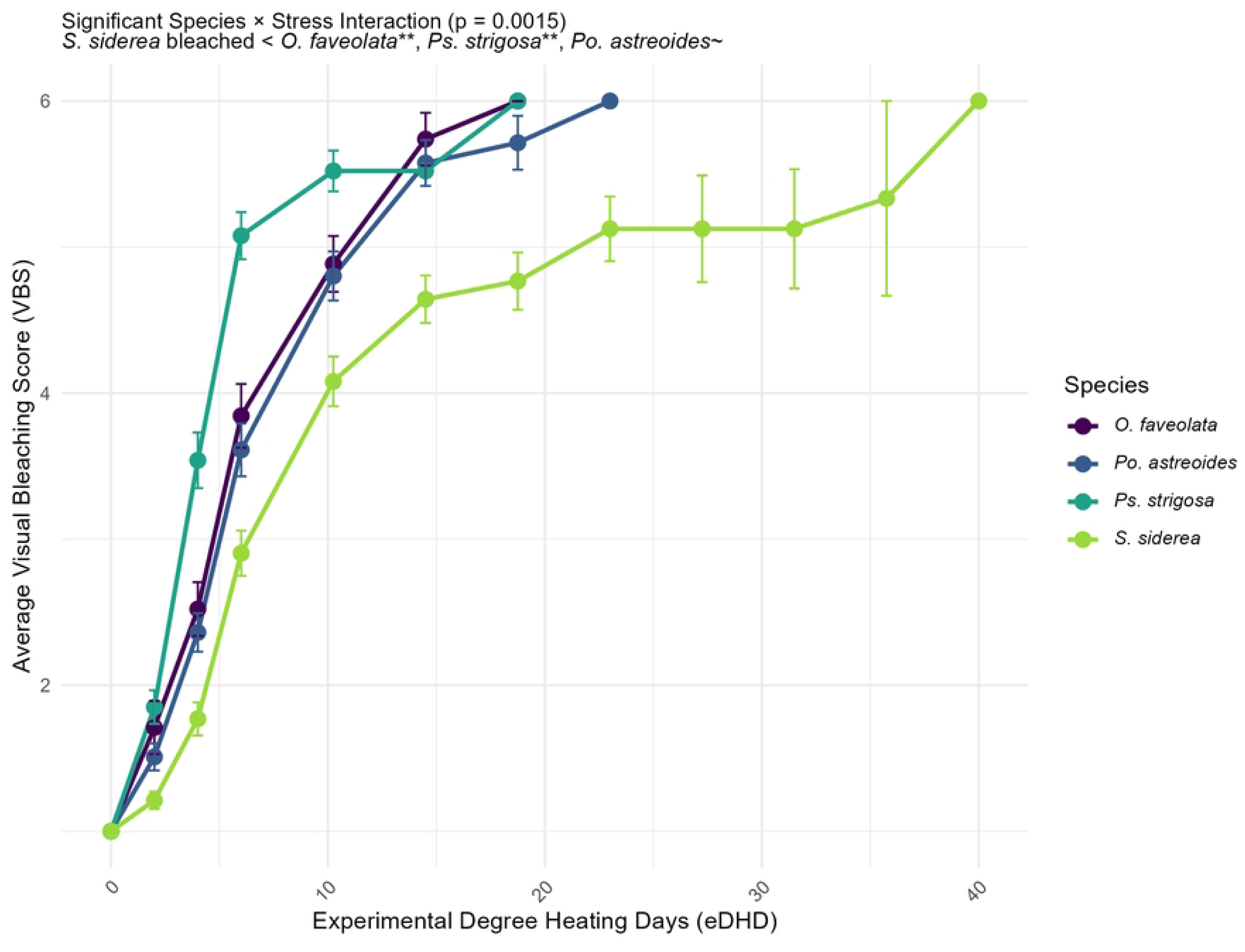
Average Visual Bleaching Responses of Belizean Coral Species by Experimental Degree Heating Days (eDHD) Progression of average Visual Bleaching Scores (VBS) for *O. faveolata*, *Po. astreoides*, *Ps. strigosa*, and *S. siderea* during heat stress experiments with increasing maximum ramp temperatures per day (°C). VBS starts at 1.0 at the baseline (T0°C) and increases through subsequent ramp temperatures, including multiple days at 37°C, capturing cumulative thermal stress effects. Error bars represent ±SE. A significant interaction was observed between species and eDHD (p = 0.0015). Based on the mean cumulative thermal stress, *S. siderea* bleached significantly less than *O. faveolata* (p = 0.0083; **), *Ps. strigosa* (p = 0.0014; **), and marginally less than *Po. astreoides* (p = 0.0529; ∼).

We next examined overall thermal tolerance, quantified as the ED50 value (the cumulative heat exposure at which individual corals appear half bleached) (Fig 2). *Siderastrea siderea* consistently exhibited the highest ED50 values among the four species across all sites, with notable peaks at 3 Corner North (11.29) and Wee Wee Caye (11.06). *Porites astreoides* displayed intermediate ED50 values, reaching a maximum at Wee Wee Caye (9.18) and decreasing to 4.63 at Carrie Bowe. *Orbicella faveolata* showed a wide range of ED50 values, from a high of 10.13 at Wee Wee Caye to a low of 3.54 at Carrie Bowe. *Pseudodiploria strigosa* had the lowest ED50 values overall, ranging from 6.36 at Bread & Butter Caye to 3.40 at Terrace. Statistical analyses confirmed these patterns, with significant interspecies differences in ED50 values (ANOVA, *p* = 2.00 × 10^−16^). Post hoc Tukey’s tests showed that *S. siderea* was significantly more resistant than all other species, with pairwise ED50 values averaging 3.83 eDHD higher than *O. faveolata* (*p* = 3.25 × 10⁻¹²), 3.03 eDHD higher than *Po. astreoides* (*p* = 1.56 × 10⁻⁸), and 5.58 eDHD higher than *Ps. strigosa* (*p* = 3.14 × 10⁻¹⁴). Additionally, *Ps. strigosa* had significantly lower ED50 values compared to *O. faveolata* (–1.75 eDHD, *p* = 0.003) and *Po. astreoides* (–2.55 eDHD, *p* = 2.77 × 10⁻⁶), while the difference between *O. faveolata* and *Po. astreoides* were not significant (*p* = 0.356).

**Fig 2.**
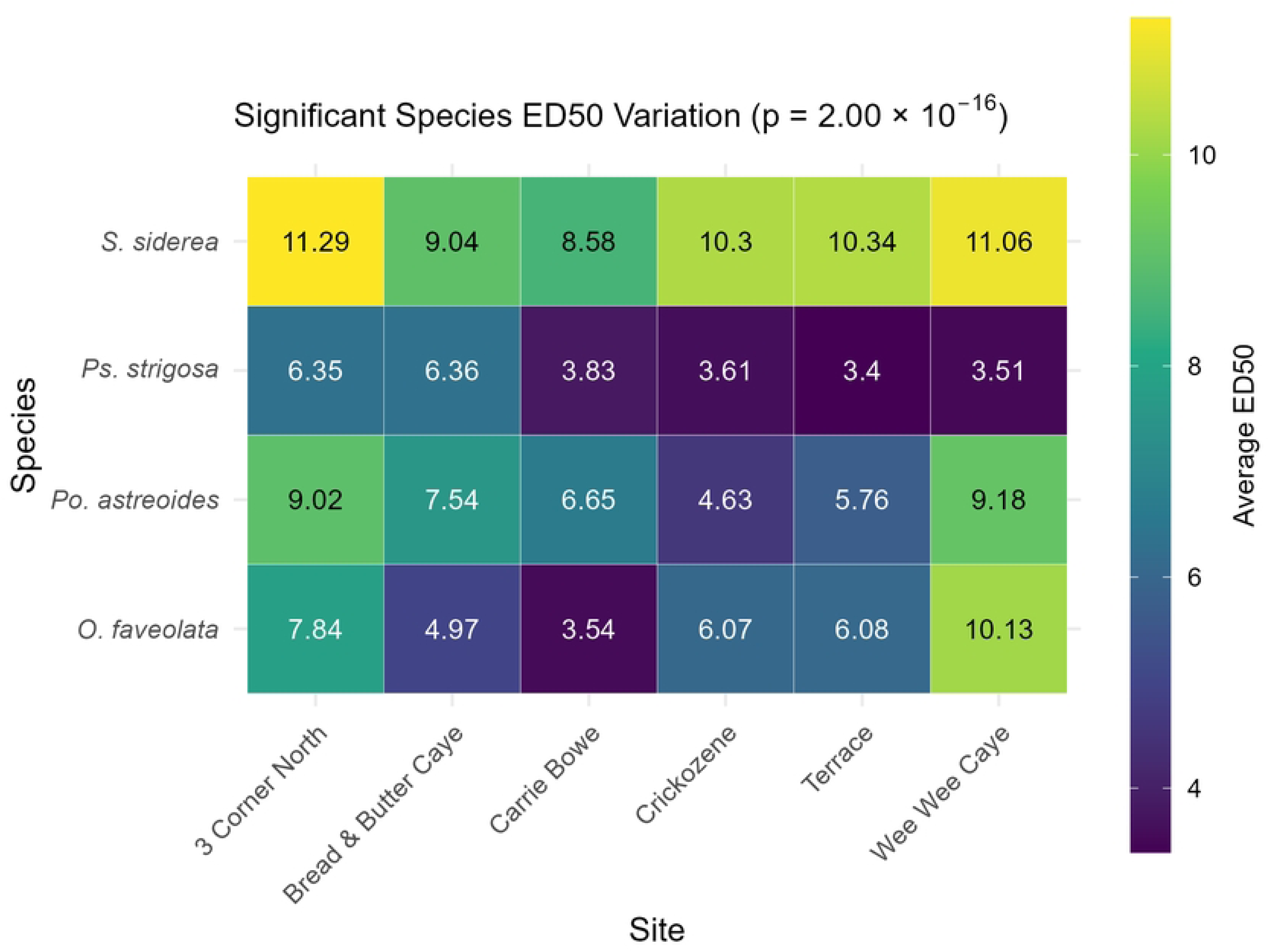
Bleaching Thresholds of Belizean Coral Species Across Sites. Heatmap showing the average ED50 values of four coral species across six Belizean reef sites. Each cell represents the average bleaching threshold (ED50 in units of DHD) for each species-site combination, with a color gradient indicating thermal tolerance ranging from low (purple) to high (yellow). ED50 marks the environmental exposure level, such as Degree Heating Days (DHD), at which coral bleaching (measured via Visual Bleaching Scores (VBS)) reaches a 50% reduction from baseline.

### Differences among locations

Site-level differences in ED50 were statistically significant (Two-Way ANOVA; *p* = 1.53 × 10⁻⁷; Fig. 3). An interaction effect between species and site locations (*p* = .001) was also identified from an ANOVA. Post hoc Tukey’s tests showed that Wee Wee Caye exhibited the highest thermal resistance for *O. faveolata* and *Po. astreoides*, with significantly greater ED50 values compared to all other sites. For *O. faveolata*, ED50 values at Wee Wee Caye were 6.59 eDHD higher than Carrie Bowe (*p* = 3.00 × 10⁻⁸), 5.17 higher than Bread & Butter Caye (*p* = 7.05 × 10⁻⁶), 4.06 higher than Crickozene (*p* = 4.47 × 10⁻⁴), and 4.05 higher than Terrace (*p* = 4.61 × 10⁻⁴). For *Po. astreoides*, Wee Wee Caye was 4.55 eDHD higher than Crickozene (*p* = 1.93 × 10⁻⁴) and 3.42 higher than Terrace (*p* = 8.22 × 10⁻³).

**Fig 3.**
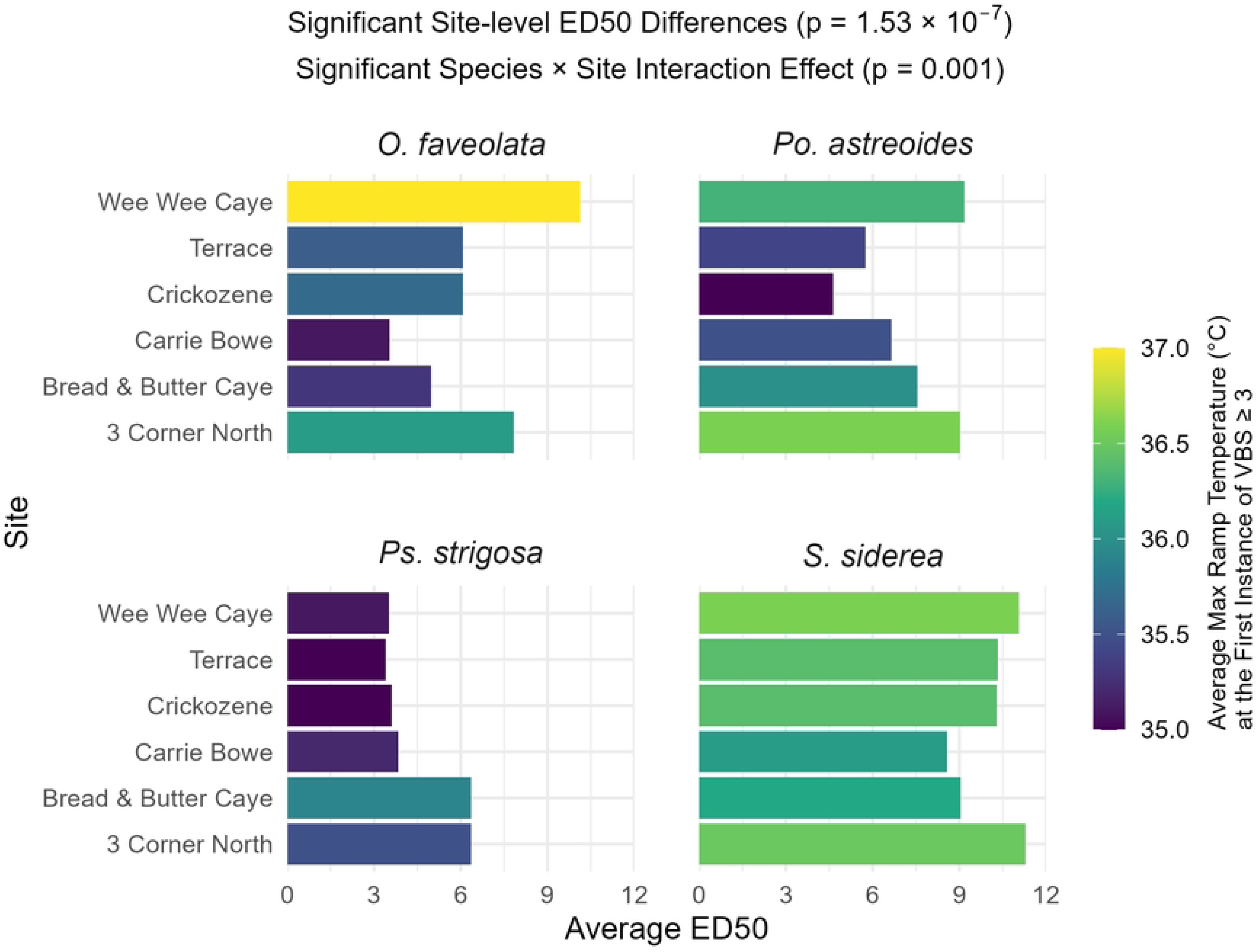
Bleaching Thresholds of Belizean Coral Species Across Sites. Average ED50 values of four coral species across six Belizean reef sites. Bar length indicates the bleaching thresholds (ED50 in eDHD), while color denotes the average max ramp temperature (°C) at which significant bleaching (VBS ≥ 3) was first observed.

3 Corner North also showed significantly elevated ED50 values. For *O. faveolata*, ED50 values at 3 Corner North were 4.29 eDHD higher than Carrie Bowe (*p* = 2.05 × 10⁻³). For *Po. astreoides*, ED50 values at 3 Corner North were 4.39 higher than Crickozene (*p* = 3.43 × 10⁻⁴) and 3.26 higher than Terrace (*p* = 1.35 × 10⁻²). For *Ps. strigosa*, 3 Corner North was significantly more resistant than all other sites. ED50 values were 2.52 to 2.95 eDHD higher than Carrie Bowe, Crickozene, Terrace, and Bread & Butter Caye (*p* = 0.003 to 1.54 × 10⁻⁴). However, *Ps. strigosa* at Wee Wee Caye had significantly lower ED50 values compared to Bread & Butter Caye and 3 Corner North (*p* = 7.08 × 10⁻⁴ and 7.44 × 10⁻⁴, respectively). Although significant site-level differences in ED50 were noted for the other species, *S. siderea*, the most heat-resistant species overall, had consistently high ED50 values, resulting in no statistically significant differences across sites.

### The top 25% most heat-resistant colonies for each species

Within species, we found marked variation in heat resistance among colonies, with the standard deviation of ED50 within each species accounting for approximately 39% of the mean (range: 35% to 44%, S1 Table). We focused on the top 25% ED50 values within species (Table 1) to estimate how much more heat-resistant corals from this upper quartile are compared to the species overall. The average ED50 of the top 25% for each species is about 53% higher than the overall average (47% to 56%), about 1.5 standard deviations above the mean (range 1 – 2 among species, Table 1). This deviation shows that a collection of the top quartile of heat-resistant corals from each species would have 1.52 DHD higher heat resistance, ranging from an increase of ED50 of 2.4 – 5.3 DHD among all species.

**Table 1.**
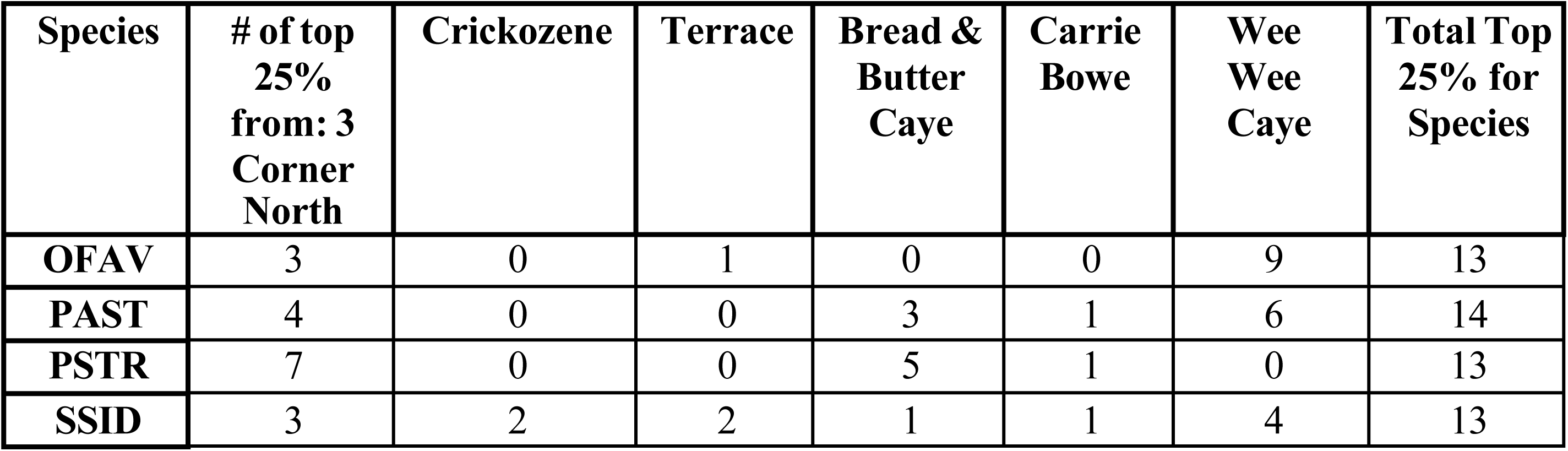
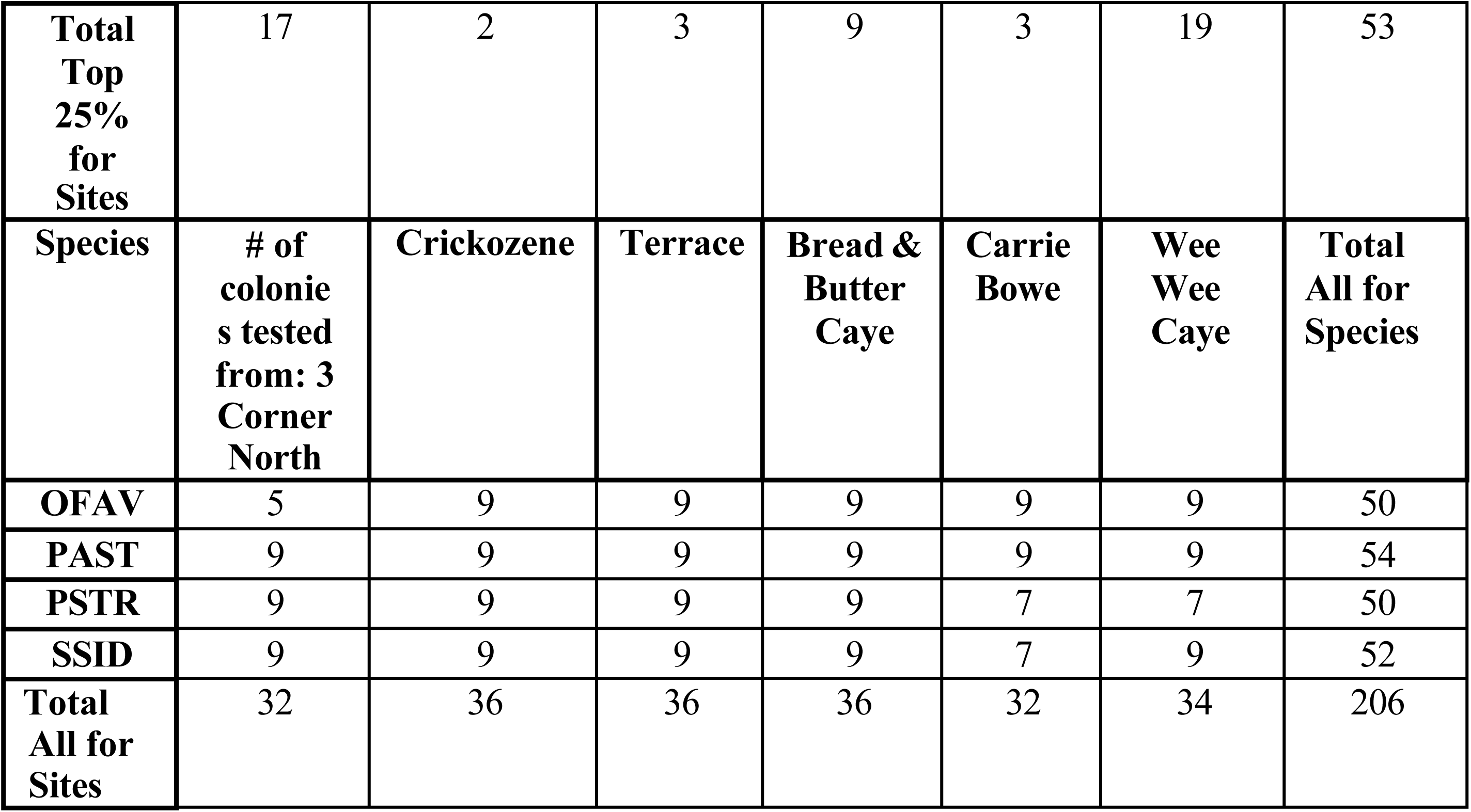
Number of coral colonies in the top 25% collected per species across six sites.

### The location of the top 25% most heat-resistant for each species

Another use of the top quartile for each coral species is to visualize where the most heat-tolerant colonies are found (Fig 4). While highly heat-resistant corals were found at every site, their distribution varied by species and location. For *O. faveolata*, 100% (9 out of 9) of the colonies sampled at Wee Wee Caye were in the top 25% of heat resistance values. This was the only site where all sampled individuals within a species fell within the top-quartile. Those nine individuals alone accounted for 62.9% of all top-quartile *O. faveolata,* with individuals from 3 Corner North contributing an additional 23.1%. A chi-squared test confirmed that the distribution of top-quartile *O. faveolata* was significantly skewed toward Wee Wee Caye and 3 Corner North (chisq test, p= 2.35×10^−5^, S2 Table).

**Fig 4.**
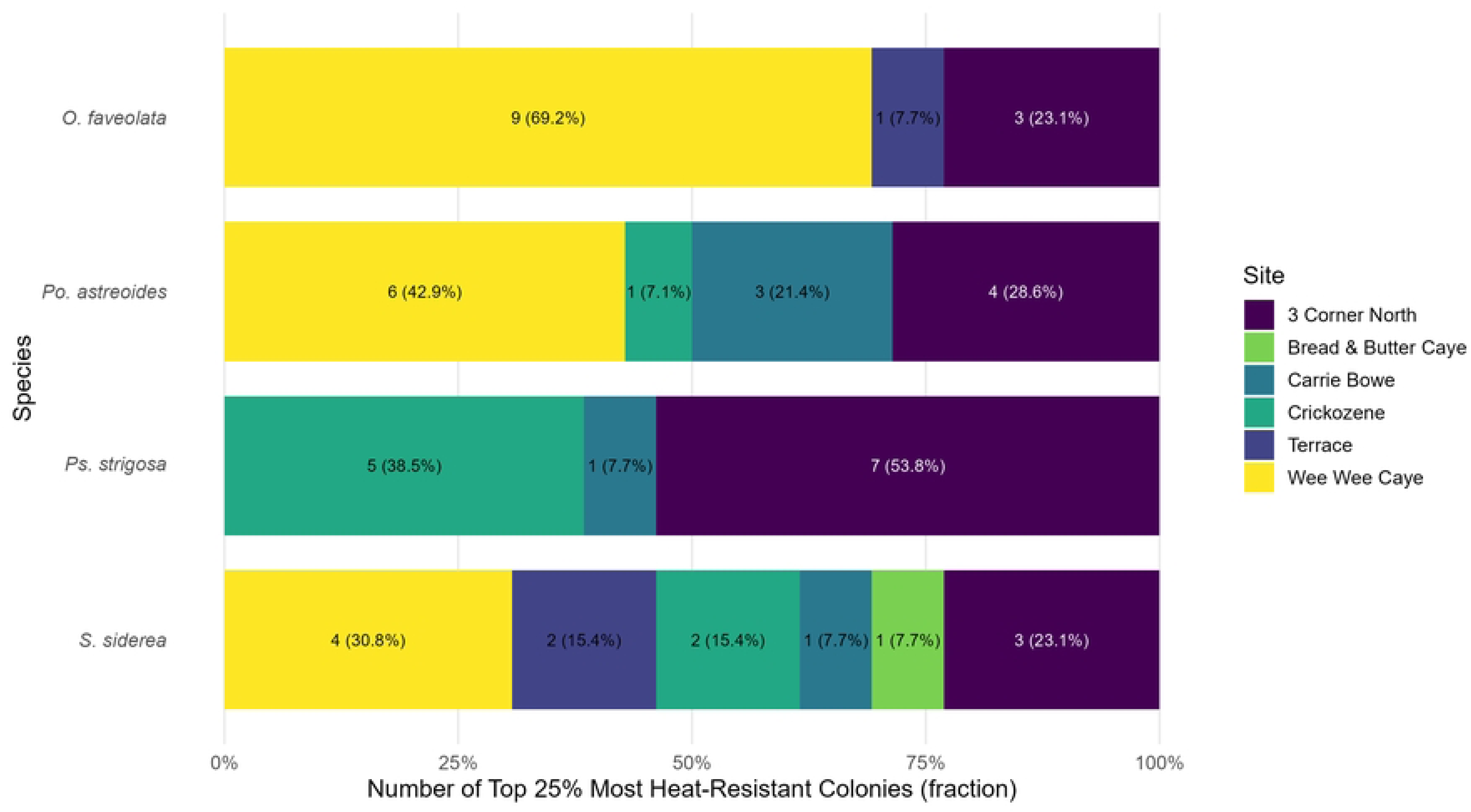
Site Contributions to the Top 25% Most Heat-Resistant Corals by Species. Each bar represents the proportion of top 25% thermally resistant corals from each site, emphasizing the dominance of certain locations in supporting high-performing colonies.

Patterns were similar for other species. We found more top-quartile *Po. astreoides* at 3 Corners North and Wee Wee Caye (chisq p=0.02), and more top-quartile *Ps. strigosa* at 3 Corner North (chisq p=0.001). In contrast, top-quartile colonies of *S. siderea* were found at similar frequencies across all sites. Although there were more top-quartile *S. siderea* found on 3 Corner North and Wee Wee Caye, this distribution was not significantly different from neutral expectation (chisq p= 0.74).

## Discussion

This study used controlled, short-term heat tolerance assays to quickly identify coral populations with natural resistance to thermal stress on six Belizean reefs. Among the four species tested, *S. siderea* was the most heat-tolerant, withstanding up to twice the experimental degree heating days (eDHD) compared to less tolerant species like *Ps. strigosa* (Fig. 2). Additionally, we found variation in heat resistance among colonies within species, similar to other findings from short-, medium-, and long-term heat stress experiments [27,29,35,36]. The greatest number of top-performing heat-resistant corals across our sites was found at the shallow-water reef at Wee Wee Caye. This site is known for high temperatures and large daily temperature variations [3], similar to previous studies that link diurnal and large daily temperature variation to increased heat resistance [29,33,37]. Coral populations at 3 Corner North, along the northern fore reef of Turneffe Atoll, also exhibited high heat-tolerance, despite the site’s likely moderate thermal conditions compared to more variable environments like Wee Wee Caye. Our study expands the use of rapid, short-term heat stress assays as an effective and scalable approach for identifying naturally heat-tolerant coral populations *in situ*, providing actionable information that can serve as a foundation for targeted conservation and restoration strategies.

### Heat Resistance Across Differing Reef Conditions

We found the highest proportions of heat-resistant corals at the shallow sites of Wee Wee Caye and 3 Corner North. All nine *O. faveolata* colonies we tested from Wee Wee Caye ranked in the top 25% for heat resistance across all 53 colonies in our study (Table 1). Overall, 19 of 34 colonies tested from this site fell within the top resistance quartile. An exception was *Ps. strigosa*, our least heat-tolerant species, which had no colonies from Wee Wee Caye in the top quartile.

Coral colonies collected within Wee Wee Caye were from a shallow patch reef located 11.74 miles from shore within the lagoon system. Long-term in situ SST records collected by Lobel & Lobel (2024) from a site farther offshore (16°45.39 N, 88°8.03 W) but within the same lagoon document sustained and rising thermal stress. Daily temperatures recorded every hour ranged from 22.3°C to 32.4°C, with a statistically significant warming trend per year over time and an increasing mean annual temperature rate of 0.0805°C per year between December 2003 and March 2014 [6]. Within this period, daily maximum temperatures frequently exceeded the regional bleaching threshold of 31.5°C during the warm season (May-September) in multiple years between 2007 and 2013.

Lagoonal habitats behind the Belize MBRS are characterized by high variability and thermal stress [3]. Baumann et al. (2016) classified these lagoonal reefs using four temperature parameters (TP): (1) yearly maximum temperature, (2) mean annual temperature range, (3) mean number of days per year that SST exceeds the regional bleaching threshold, and (4) mean length of the longest consecutive run of days above that threshold. Sites classified as High_TP_ sites are reefs that have elevated values across all four-parameter metrics, with average mean annual maximum temperatures ranging from 30.9 to 31.3°C, annual temperature ranges of 5.9 to 7.1°C, 54.4 to 78.4 annual days above the bleaching threshold, and 5.7 to 7.5 consecutive days above the bleaching threshold. These environmental conditions likely create localized thermal stress regimes that may contribute to the maintenance or selection of heat-tolerant coral populations [3,6,24,27,33,38–40].

While identifying High_TP_ environments has been useful for locating reefs likely to have thermally resistant corals, our findings also show that heat-resistant populations are not exclusive to these environments. Most of Turneffe Atoll, excluding the central lagoon, is classified as a low_TP_ region due to cooler average temperatures and fewer days above the bleaching threshold [3]. However, the back reef at 3 Corner North, a shallow reef flat behind the reef crest of northern Turneffe Atoll, is not included in these classifications due to gaps in available data. Unlike the sheltered central lagoon, which experiences reduced wave energy and limited tidal flushing [34], the northern section of Turneffe atoll is exposed to greater wave action and open circulation from the Caribbean [41,42]. Such conditions are associated with stronger tidal action flushing, more stable temperatures, and reduced chronic thermal stress [43,44], and are therefore expected to support less thermally preconditioned coral populations.

Yet, at 3 Corner North, we recorded unexpected high heat resistance across species, particularly in *Ps. strigosa*, the least heat-tolerant species overall in our study. Of the nine *Ps. strigosa* colonies tested from this site, seven ranked among the most heat-tolerant (Table 1), and on average, these colonies were twice as heat-resistant as same-species colonies from other locations (Fig. 3). Similar patterns of high heat resistance found in moderate-temperature environments were reported by Cornwell et al. (2021), who documented high heat tolerance at a fore reef site in Palau. Together, these results suggest that high heat tolerance can emerge under various environmental conditions.

### Species Variation in Heat Tolerance

Our results show that *S. siderea* demonstrated consistently higher thermal resistance than the other three species across all six sites tested. This aligns with findings by Baumann et al. (2016), who documented this species’ success throughout Belize’s thermally variable reefs, including high-temperature sites. This species possesses structural traits such as slow growth, massive morphology, and long lifespan, supporting its classification as stress-tolerant [38], and it can maintain positive calcification under elevated temperatures and acidification [45,46]. Nearshore *S. siderea* were also shown to host *Durusdinium trenchii,* an algal symbiont with higher thermal tolerance [47]. Together, these structural, physiological, and symbiotic features may help explain the species’ broad ecological success across various thermal regimes, which may position *S. siderea* as a key survivor on future reefs under climate change [3,38].

Across the locations we tested, *Ps. strigosa* consistently exhibited the lowest heat resistance of all species in our study. Although both *S. siderea* and *Ps. strigosa* are classified as stress-tolerant species by Baumann et al. (2016), subsequent studies suggest that *Ps. strigosa* is more physiologically vulnerable. For example, Bove et al. (2022) list *Ps. strigosa* as sensitive to heat and elevated CO_2_ [46], and Aichelman et al. (2021) observed progressive physiological deterioration in this species under prolonged warming [48]. In Belize, *Ps. strigosa* hosts a variety of symbionts across thermal environments, including the less heat-tolerant *Breviolum minutum*, even on reefs with moderate and high temperatures. Some colonies also host various strains of *Cladocopium,* and a smaller proportion of algal cells have been identified as *Durusdinium* [49]. This symbiont diversity, particularly the dominance of thermally sensitive taxa, may influence the species’ vulnerability to heat stress.

Unlike the consistent response to heat stress across sites from *S. siderea* and *Ps. strigosa,* thermal tolerance for *Po. astreoides* and *O. faveolata* varied by location. Site-level variability was most pronounced in *O. faveolata,* a generalist species with moderate growth, structural complexity, and intermediate stress tolerance [38]. Baumann et al. (2016) reported few *O. faveolata* colonies at high-temperature sites in Belize; yet, our data show high performance of this species at one of the warmest and most variable sites in our study. All nine colonies sampled from Wee Wee Caye ranked among the most heat-resistant individuals, even outperforming *S. siderea*. In contrast, colonies from nearby reefs such as Carrie Bow Caye and Bread and Butter exhibited only half this level of heat resistance (Fig. 3).

Thermal tolerance in *O. faveolata* has been shown to depend on a combination of factors, including reef position, symbiont composition, host genetics, and acclimation. Bleaching resistance in this species has been linked to the presence of the heat-tolerant symbiont *D. trenchii* and distinct microbial communities at a 33 °C reef in the Bahamas [50]. Similarly, symbiont identity and host genotype largely explained bleaching resistance in the Florida Keys [51]. Recent work by Aguilar et al. (2024) further supports these patterns, linking inshore thermal tolerance to host gene regulation and symbiont identity [52], while Ricaurte et al. (2024) found seasonal proteomic shifts that reflect increased physiological stress resilience [53].

Together, these findings suggest that local adaptation or acclimatization may enable some *O. faveolata* populations to exceed expected thermal limits. However, the relationship between symbiont community structure and reef position remains complex. For example, Manzello et al. (2019) found that most *O. faveolata* colonies sampled in Florida hosted multiple symbiont genera, and these did not consistently correlate to onshore/offshore positions [51]. Aguilar et al. (2024) further demonstrated that symbiont type can shift with incubation at high temperatures, with offshore colonies tending to host *Cladocopium,* whereas inshore populations had a variety of genera. These symbiont differences were also influenced by host genotype, suggesting that interactions between host genetics and symbiont identity contribute to differential bleaching susceptibility [51].

Unlike the slow-growing, stress-sensitive *Orbicella* corals, *Po. astreoides* is an opportunistic species predicted to persist under increasing thermal stress due to its rapid growth, high fecundity, and flexible recruitment strategies [38]. Consistent with this expectation, *Po. astreoides* have been observed to dominate many high-temperature reef sites in Belize [3,54]. The brooding and parthenogenetic reproduction of *Po. astreoides* likely contributes to the prevalence of this species at high-temperature sites [55]. Yet, our data suggests that this weedy life history alone does not fully explain the species’ success. In Florida, *Po. astreoides* experienced higher natural bleaching in regions with greater heat stress [56], a pattern not linked to symbiont identity. In contrast, our results show that *Po. astreoides* exhibited the second-highest heat resistance of all species tested, with high ED50 values recorded at Wee Wee Caye and Three Corners North. These findings point to elevated physiological thermal tolerance in addition to its opportunistic traits.

The variation in thermal resistance observed across species and sites highlights the utility of conducting population-specific assessments to identify corals with elevated heat tolerance. While life history traits provide a useful framework, our results demonstrate that bleaching susceptibility often reflects a combination of species-level traits, local environmental conditions, and population-level differences. Multiple mechanisms are increasingly recognized as contributors to coral heat resistance, including host genotype and gene expression [40,57], symbiont community composition [27,49], and the coral microbiome [58]. However, a deeper understanding of how these factors interact remains an active and important area of research.

### Using Rapid Test of Coral Heat Resistance in Conservation Management

Even in the face of complex mechanisms underlying coral heat resistance, using a standardized, reliable, and cost-effective framework for identifying naturally heat-tolerant populations makes species- and location-specific discoveries feasible for targeted reef protection and restoration. Our experimental setup, which relies on simple equipment and minimal infrastructure (e.g. Palumbi et al. 2014), serves as the foundation for more complex systems such as CBASS, used in recent experiments [59]. Our version provides an inexpensive and portable platform for testing corals across species and locations, enabling scalable assessments of thermal tolerance in local reef environments. This approach supports conservation and restoration by helping prioritize coral populations most likely to persist under future thermal stress.

From this framework, two complementary approaches emerge for increasing reef resistance under climate change [60]. First, conservation efforts could prioritize protecting reef sites that harbor a large population of heat-resistant corals, helping to preserve traits that enable natural thermal tolerance. Such an approach does not depend on the mechanisms of heat tolerance if resistance is stable at the site level. Protecting these heat-resistant sites aligns with national goals under the Resilient Bold Belize initiative, which aims to expand fully protected coral reef areas from 7% to 20% by 2026.

Second, restoration programs can use thermally resistant populations as donor stock for selective propagation and assisted migration to areas that may provide refuge and benefit from enhanced resistance. For example, using locally sourced, heat-resistant colonies in coral nurseries has been shown to improve bleaching outcomes across four species in American Samoa during natural heatwaves [32]. Models of local coral adaptation also suggest that increasing the number of reproductive colonies with heat-resistant corals can facilitate faster adaptation to future climate conditions through assisted evolution [57,61–63].

The success of these strategies depends on the durability of thermal resistance after transplantation and whether laboratory-based assessments can reliably predict performance during field-based heatwaves [32,64,65]. Such studies are likely to be species-, location-, and symbiont-dependent and can be accomplished with common garden experiments, using recent restoration techniques and coral re-testing with the kind of portable lab setup we employed here.

Protecting and utilizing thermally resistant populations not only sustains ecological function under warming conditions but also increases the likelihood of long-term success of restoration efforts. These populations also provide opportunities to supply heat-resistant colonies of many species to restoration nurseries, helping address the dual need for increased thermal tolerance and biodiversity. This highlights the essential role of functional diversity in maintaining reef resistance and ecological integrity in a changing climate.

## Conclusion

This study identified coral species and populations across Belizean reefs that exhibit natural resistance to elevated temperatures. Heat-tolerant colonies were found at all sites, with notable differences in ED50 values among species and locations. The variation in heat-tolerance observed across and within species and reef sites highlights the complex interplay of biological and environmental factors that shape coral thermal tolerance. While the underlying mechanisms remain an active area of research, the ability to identify naturally heat-tolerant corals using inexpensive and reliable testing provides an immediate pathway to support restoration and conservation. These populations also offer valuable opportunities for ongoing investigation into the factors that drive coral bleaching resistance.

## Materials and Methods

### Characterizing Belizean Coral Heat Tolerance

Identifying corals with elevated heat resistance can inform national priorities for marine protection and restoration. However, the 190-mile span of Belize’s MBRS and various reef locations created logistical challenges for identifying priority sites. To overcome these challenges, we considered factors like target species coverage, testing setup location, transport conditions, and local temperature. Therefore, we focused on four coral species: *Siderastrea siderea*, *Porites astreoides*, *Orbicella faveolata*, and *Pseudodiploria strigosa* across three locations within Turneffe Atoll and South Water Caye Marine Reserve (SWCMR).

We used environmental maps to provisionally identify possible areas for future protection with high coral cover or unique features like increased nutrient enrichment or high temperatures. Sea surface temperature (SST) and preliminary heat helped refine target area selections, focusing on capturing temperature variability across sites. Atlantic and Gulf Rapid Reef Assessment (AGRRA) survey data, gathered before the 2023 bleaching event, directed us towards reefs experiencing various temperatures, had a reef health index score of 2.8 or higher, and were at similar depths to reduce environmental variability.

### Site Evaluation

The sites selected for heat tolerance testing met specific criteria: all target species must be present to ensure consistent sampling, sites should be at snorkel depth for accessibility and to limit environmental variance inconsistency, and locations needed to be within an hour of the heat stress tank location to minimize coral stress during transport. Sites were required to span a wide temperature range to capture potential phenotypic responses to warming, allowing for the identification of heat-tolerant individuals and reefs.

While the warm temperatures near Placencia made it a region of interest, an accessible location for conducting experiments was difficult to find. The same issue arose near Belize City, although St. George’s Caye holds potential for future research. With these two areas omitted, we focused on sites near Tobacco Caye and Glovers Atoll. SST temperature data also led us to prioritize Turneffe Atoll, known for its cooler water, despite its initial exclusion due to its protected status. Ultimately, three sites from Turneffe Atoll (3 Corner North, Terrace, and Crickozene) and Tobacco Caye (Carrie Bow Caye, Wee Wee Caye, and Bread & Butter) were chosen for the experiments (Fig 5). Before starting the experiments, an additional scouting trip enabled us to select specific sites within Turneffe Atoll.

**Fig 5.**
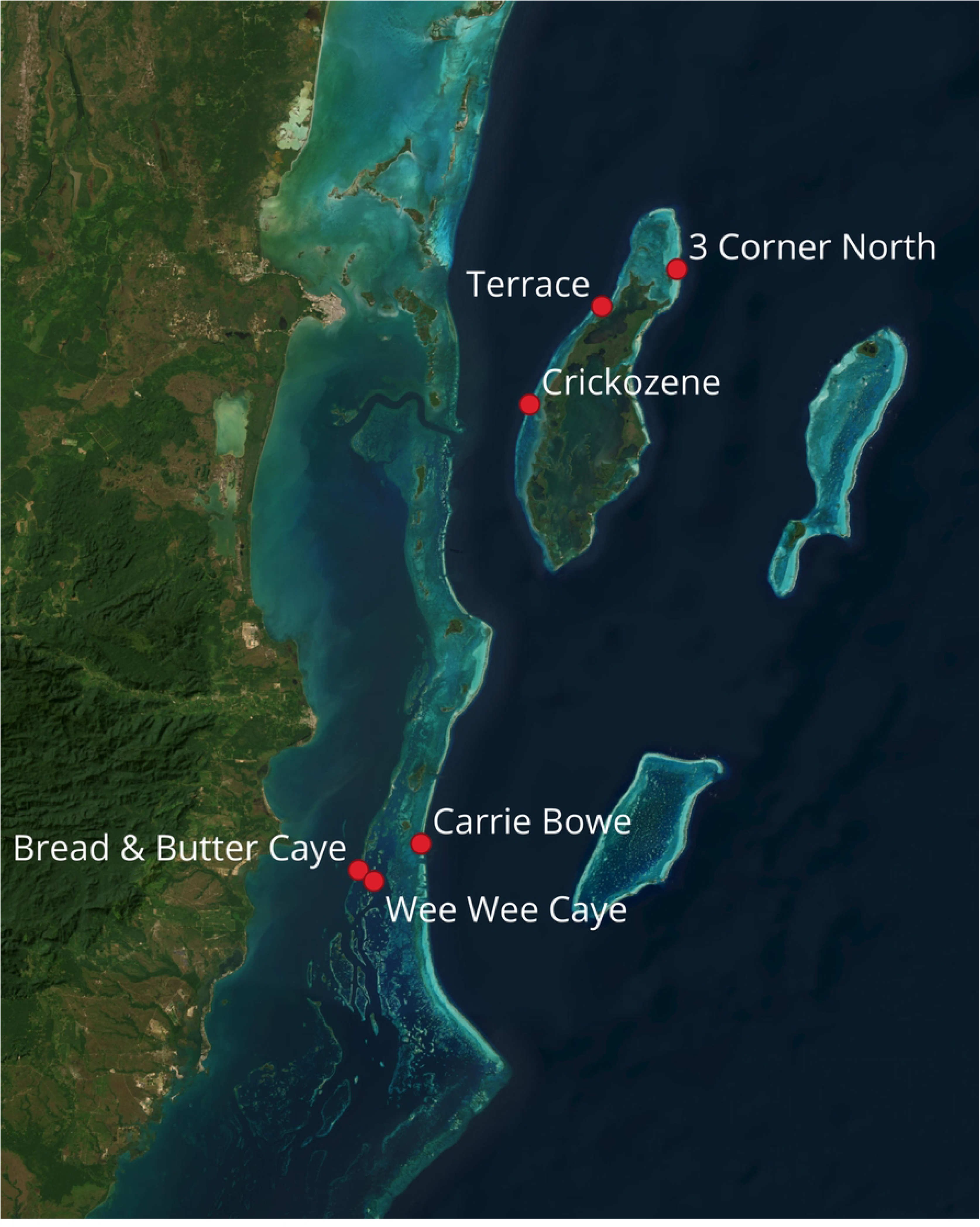
Reef sites prioritized for coral heat tolerance studies across Belize. The selected locations—3 Corner North, Terrace, Crickozene, Bread & Butter, Carrie Bow, and Wee Wee—were chosen based on coral diversity, colony abundance, and accessibility. These sites represent key areas for testing and studying thermal resistance, encompassing diverse reef ecosystems and conditions. Their strategic selection ensures comprehensive coverage for effective heat stress experiments.

### Coral Collection

Multiple-day short-term heat experiments were conducted at the University of Belize Calabash Caye Field Station (UBCCFS) and the Tobacco Caye Marine Station (TCMS) within SWCMR. Three multi-temperature experimental series were conducted at each base, with each series corresponding to a different collection reef. In total, six series were completed, three for each of the two selected reef regions.

For each experimental run, 5-9 replicate fragments (5-7 cm^2^) were taken from *O. faveolata*, *Po. astreoides*, *Ps. strigosa*, and *S. siderea* colonies using hammers and chisels at each site. Each colony was tagged with a unique numbered cow tag, and its location was recorded using a handheld GPS on the surface (Garmin, Lenexa, KS). On April 22^nd^, 2024, five *O. faveolata* colonies and nine *P. astreoides*, *P. strigosa*, and *S. siderea* were collected randomly from healthy colonies at 3 Corner North at (∼1 m depth). Only five *O. faveolata* colonies were collected because four were later identified as *Orbicella annularis*. Additional collections were made at Crickozene on April 26^th^ and Terrace on April 30^th^. Nine *O. faveolata*, *Po. astreoides*, *Ps. strigosa*, and *S. siderea* colonies were collected at depths ranging from ∼1.5 m to ∼2.7 m. In SWCMR, nine colonies of each species were collected from Bread and Butter (∼2.2 m) on May 10^th^,2024, Carrie Bow (∼1.8 m) on May 14^th^, and from Wee Wee Caye (∼1.6 m) on May 18^th^. A total of 206 coral ramets from the six sites were collected for heat tolerance testing **(Table 3)**.

**Table 3:** Coral species fragments collected at the final six selected sites for heat tolerance studies. The columns represent the six selected sites. The rows show the number of fragments collected for each target species: O. faveolata, P. astreoides, P. strigosa, and S. siderea. Total All for Sites: Indicates the total number of fragments collected at each site.

| Species | 3 Corner North | Crickozene | Terrace | Bread & Butter Caye | Carrie Bowe | Wee Wee Caye | Total for all Species |
| --- | --- | --- | --- | --- | --- | --- | --- |
| *O. faveolata* | 5 | 9 | 9 | 9 | 9 | 9 | 50 |
| *Po. astreoides* | 9 | 9 | 9 | 9 | 9 | 9 | 54 |
| *Ps. strigosa* | 9 | 9 | 9 | 9 | 7 | 7 | 50 |
| *S. siderea* | 9 | 9 | 9 | 9 | 7 | 9 | 52 |
| **Total**<br>**All for**<br>**Sites** | 32 | 36 | 36 | 36 | 32 | 34 | 206 |
| **Coordi**<br>**nates** | 17°32'59.<br>06"N<br>87°45'5.2<br>8"W | 17°22'29.02"<br>N<br>87°56'32.77"<br>W | 17°30'6.0<br>2"N<br>87°50'55.<br>50"W | 16°46'17.98"N<br>88° 9'50.54"W | 16°48'21.<br>49"N<br>88°<br>4'59.27"<br>W | 16°45'26.<br>64"N<br>88°<br>8'39.12"<br>W |  |

Coral fragments from the collection were stored in storage baskets divided into six bins. Each compartment had a numbered tag to distinguish between collected colonies. After collecting all species, the baskets were transported back to the base in seawater-filled covered containers with regular water changes every 20 minutes. Once back, the fragments were further divided into smaller ramets and then placed in open containers in shallow ocean water near the shore to recover overnight.

### Stress Tank Protocol

From April 24 to May 25, 2024, six acute heat experiments were conducted using five 48 Qt experimental tanks. Each tank temperature was regulated using a 200 W heater connected to an Inkbird (ITC-608T) programmed to run one of two thermal profiles, control or heat. Tanks always had fresh seawater inflow from spigots at 16 mL/10 seconds or ∼one full volume change every 7 hr and two aquarium pumps (∼280 L hr–1) to increase flow around the fragments and circulate temperatures. The inflow was adjusted to 25 ml/10 seconds during the approximately three-hour cooling period of the experiments. Natural light with a shade cloth (∼100-300 PAR) illuminated the tanks. To monitor the thermal environment of the tanks, we attached two HOBO temperature loggers (Onset Computing, Bourne, MA) to each coral rack and set to record temperature at 10-minute intervals.

In the morning after overnight sampling recovery, one coral ramet per colony per species (n=36) was randomly placed within a designated species row within a tank with a starting temp of ∼29°C. Two tanks served as the control treatments, starting at ∼29°C and increasing by 1°C daily until 32 °C. Two tanks were subjected to experimental heat ramps starting at 9:30 am: temperatures ramp up (increase) to the respective target maximum temperature over three hours (1100-1400), followed by a three-hour hold at maximum temperature (1400-1700), a 4-hour ramp down in temperature to 30°C (1700-2100), and an overnight “recovery” at 30°C. The following morning, at 9:30 am, the heat ramp temperatures increased by 1°C. This increase in temperature would continue for four days until the experimental tank reached a maximum temperature of 37°C and recovery temperature of 32°C. On the fifth day, any surviving coral ramets from the two experimental tanks were randomly subsampled to represent that colony in a fifth tank that continued the heat ramp program starting at 32°C, reaching a maximum temperature of 37°C, and resting again at 32°C.

We visually evaluated coral fragments at 8:00 AM each day of the experiment (one observation before each ramp). We visually assessed coral fragments using a five-point scale that consisted of the following categories: 1 – no bleaching, 2 – slight bleaching visible, 3 – moderately discolored, clearly bleaching, 4 – severe bleaching, with some color remaining, 5 – no color, completely bleached. We evaluated each fragment using four scorers who were required to agree on a score for each fragment during each assessment.

### DHD and ED50 Calculation

To quantify the upper thermal thresholds of corals, experimental Degree Heating Days (DHD) [67] were calculated as a measure of cumulative thermal stress experienced during the acute thermal assays. This calculation incorporates ramp and night stress into Degree Heating Hours (DHHR), which represents the accumulation of heat-loading hours over the regional bleaching threshold [35]. Together, DHHR values form a continuous stress framework, starting with an initial baseline temperature (*Tavg local temp*) and dividing by 24 to calculate DHD.

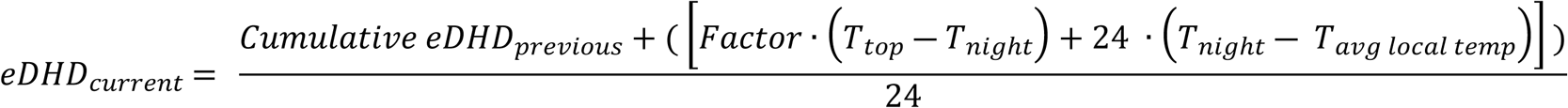

Thermal ramp cumulative stress is determined by multiplying a scaling factor (6), representing the duration of the ramp phase relative to the total daily thermal stress, with the difference between the maximum ramp temperature (Ttop) and the nighttime temperature (Tnight). The night stress is calculated as the product of 24 hrs and the difference between Tnight and the average local temperature (Tavg local temp). These stress contributions are summed daily and added to the (DHHR from previous days) divided by 24 to yield the eDHD. The cumulative approach reflects the progressive thermal load experienced by corals. The eDHD values are used to determine ED50, which represents the DHD level at which corals reach a VBS of 3. ED50 is estimated as the DHD at which the best linear regression of VBS against DHD 3.0 [ED50= (3-intercept)/slope]. This approach provides a quantitative threshold for coral thermal tolerance under experimental or natural conditions, which is instrumental in comparting thermal resistance across sites.

### Statistical Analysis

To evaluate the effects of coral species, site, and their interaction on thermal tolerance measured by ED50 values, a two-way analysis of variance (ANOVA) was performed. Tukey’s Honest Significant Difference (HSD) test was also applied to perform pairwise comparisons on any significant effect. A chi-squared test was conducted to evaluate whether the distribution of the most heat-resistant coral colonies (those in the top 25% of heat resistance) varied significantly across sites for each species. Post-hoc analyses were performed to identify specific sites with significant deviations, using standardized residuals to quantify the magnitude of differences between observed and expected counts. This approach allowed for a detailed assessment of spatial patterns in coral heat resistance, highlighting sites that disproportionately contributed to the population of top-performing colonies.

## Acknowledgements

The authors would like to acknowledge the staff and boat operators of the UBCCFS and TCMS. Furthermore, the authors would like to thank the Belize Fisheries Department for their support of this project. Basemap imagery provided by Esri, Maxar, Earthstar Geographics, and the GIS User Community via ArcGIS World Imagery (EPSG:3857). Map created using QGIS.

## Supporting Information

**S1 Table.** This table provides a statistical overview of ED50 values for *O. faveolata, Pa. astreoides, Ps. strigosa, and S. siderea,* based on their average ED50, variability, and the performance of the top 25% of colonies.

| Species | avg_ED<br>50 | STD<br>std_dev | SD_mean_<br>ratio | top_25_avg_<br>ED50 | diff_ed<br>50 | top_25_sd_above_<br>mean | top_25_all_<br>ratio |
| --- | --- | --- | --- | --- | --- | --- | --- |
| **OFAV** | 6.33 | 2.79 | 0.44 | 9.89 | 3.57 | 1.32 | 1.56 |
| **PAST** | 7.13 | 2.52 | 0.35 | 10.5 | 3.35 | 1.60 | 1.47 |
| **PSTR** | 4.58 | 1.81 | 0.39 | 7.03 | 2.46 | 0.97 | 1.54 |
| **SSID** | 10.2 | 3.86 | 0.38 | 15.5 | 5.38 | 2.23 | 1.53 |
| **Average** |  |  | 0.39 | 10.74 |  | 1.53 | 1.52 |

**S2 Table.**
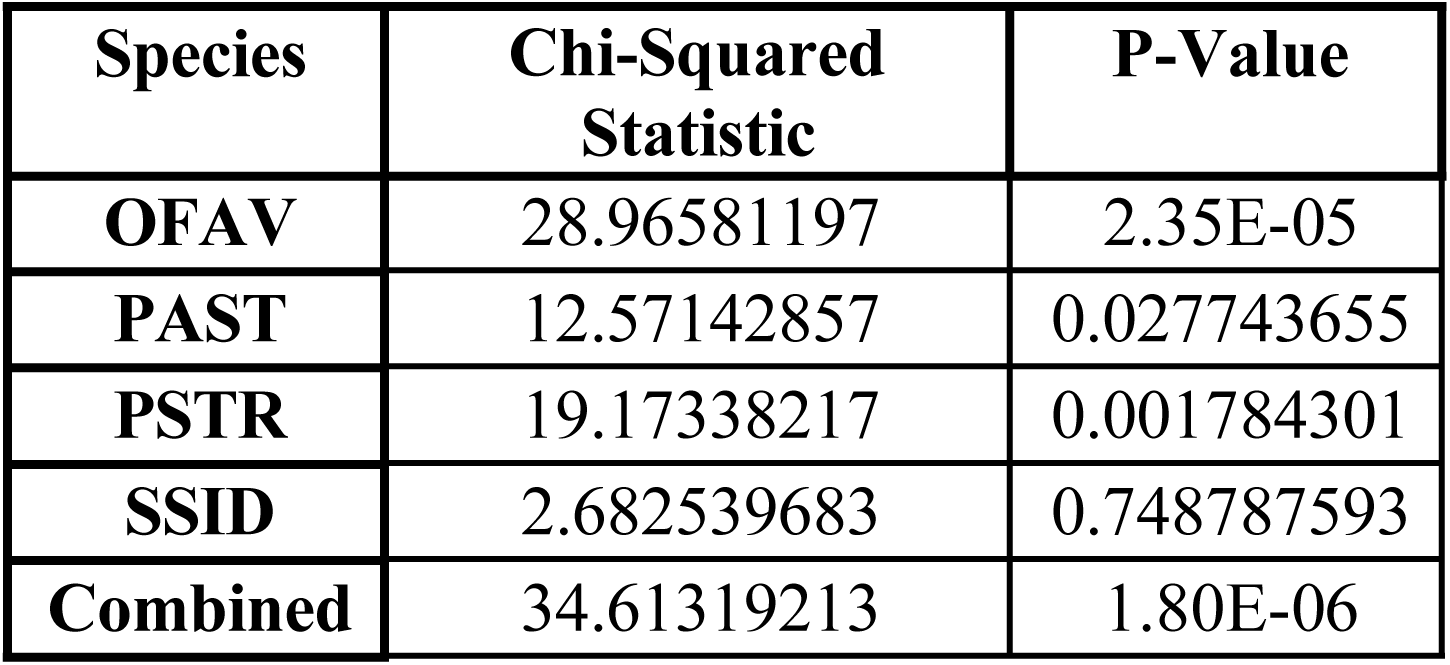
The results of the chi-squared for the distribution of thermally resilient coral colonies across sites for most species.

| Species | Chi-Squared<br>Statistic | P-Value |
| --- | --- | --- |
| **OFAV** | 28.96581197 | 2.35E-05 |
| **PAST** | 12.57142857 | 0.027743655 |
| **PSTR** | 19.17338217 | 0.001784301 |
| **SSID** | 2.682539683 | 0.748787593 |
| **Combined** | 34.61319213 | 1.80E-06 |

